# Microbial microdroplet culture system (MMC): an integrated platform for automated, high–throughput microbial cultivation and adaptive evolution

**DOI:** 10.1101/2019.12.19.883561

**Authors:** Xingjin Jian, Xiaojie Guo, Jia Wang, Zheng Lin Tan, Xin-hui Xing, Liyan Wang, Chong Zhang

## Abstract

Conventional microbial cell cultivation techniques are typically labor intensive, low throughput, and poor parallelization, rendering them inefficient. The development of automated, modular microbial cell micro–cultivation systems, particularly those employing droplet microfluidics, has gained attention for their high–throughput, parallel and highly efficient cultivation capabilities. Here, we report the development of a microbial microdroplet culture system (MMC), which is an integrated platform for automated, high–throughput cultivation and adaptive evolution of microorganisms. We demonstrated that the MMC yielded both accurate and reproducible results for the manipulation and detection of droplets. The superior performance of MMC for microbial cell cultivation was validated by comparing the growth curves of six microbial strains grown in MMC, conventional shake flasks or well plates. The highest incipient growth rate for all six microbial cell lines was achieved using MMC. We also conducted an 18–day process of adaptive evolution of a methanol–essential *Escherichia coli* strain in MMC and obtained two strains exhibiting higher growth rates compared with the parent strain. Our study demonstrates the power of MMC to provide an efficient and reliable approach for automated, high–throughput microbial cultivation and adaptive evolution.

## INTRODUCTION

The cultivation of microbes is the foundation of microbiological research and is absolutely necessary for microbial isolation (Kim et al., 2019; MaymoGatell et al., 1997), identification (Wadlin et al., 1999), screening (Lu et al., 2018; Nautiyal 1999) and evolution (Cubas–Cano et al., 2019; Dunham et al., 2002; Gao et al., 2017). Conventional, reliable laboratory cultivation techniques include shake flasks, well plates and solid medium. However, technological advances in applied microbiology, for instance, using the metabolic capacity of microbes to produce chemicals (Nielsen et al., 2013), fuels (Hong et al., 2012), pharmaceuticals (Ferrer–Miralles et al., 2009), and enzymes (Li et al., 2015), necessitate the development of cultivation methods that provide more information on strain and process properties and higher throughput than conventional methods (Hemmerich et al., 2018). Improved cultivation methods would also address limitations of conventional methods, including throughput, reagent and labor cost, and cultivation properties such as mixing and parallelization.

Recent advances in miniaturization and parallelization of microbial cultivation systems have translated into advances in the capabilities of microbioreactors, which are now used in studies of microbial antibiotic resistance and evolution (Kaminski et al., 2016; Toprak et al., 2012 & 2013), high–throughput analysis and sorting of microbes (Agresti et al., 2010; Beneyton et al., 2017; Sjostrom et al., 2014), microbial enrichment (Akselband et al., 2006; Bachmann et al., 2013), and discovery of new microorganisms (Nichols et al., 2010; Park et al., 2011). Precise control of cultivation parameters (e.g., temperature, pH and nutrient concentrations) and methods of detection (e.g., optical density, fluorescence intensity, pH and dissolved oxygen) has been achieved by implementing novel analytical technologies (Faassen & Hitzmann, 2015; Marose et al., 1999). In addition, automated operations have reduced human error in experiments, and parallelization has been achieved through miniaturization and the use of independent control systems for each microbioreactor.

Micro–cultivation systems can be classified into single–phase microfluidic cultivation systems and droplet–based cultivation systems (Tan et al., 2019a). A single–phase microfluidic cultivation system is a millimeter–scale, automatic, continuous cultivation system that is, essentially a smaller version of a conventional continuous cultivation system, e.g., a chemostat or turbidostat. These systems are usually modular and open–sourced, enabling further modification based on experimental needs (Takahashi et al., 2015; Wong et al., 2018). On the other hand, development of the micro total analysis system (commonly known as μTAS; Angell et al., 1983; Terry et al., 1979) has enabled the microscale manipulation of fluids and provided several advantages over microfluidic–based cultivation systems, e.g., high inside mass transfer rate (Lee et al., 2011) and large specific surface area, facilitating gaseous exchange (Li et al., 2012) and higher throughput. Although various microfluidic– based cultivation systems have been constructed (Balagaddé et al., 2005; Lee et al., 2011), single–phase microfluidic systems are prone to microbial contamination (Zhang & Xing, 2007). Microdroplet technology effectively addresses this problem by compartmentalization, i.e., by encapsulating microbial cells in droplets on the scale of microliters, nanoliters, or even picoliters. By separating carrier fluid from the culture medium, contamination is essentially eliminated. Microdroplet technology also has the potential for very high–throughput cultivation, which can reach hundreds (Jakiela et al., 2013; Wink et al., 2018), thousands (Baraban et al., 2011; Ota et al., 2019), or even millions of droplets in every experiment (Kaushik et al., 2017). In addition, further miniaturization of a cultivation system enhances parallelization of cultivation. For example, high–throughput, continuous cultivation of *Escherichia coli* and *Pseudomonas fluorescens* to a final cell density of 10^6^ cells per droplet has been achieved in Teflon tubes using 60–nl droplets (Grodrian et al., 2004). In addition, continuous monitoring and cultivation, adaptive evolution of microbial cells, and screening for antibiotic toxicity have been achieved in automated microdroplet cultivation systems (Churski et al., 2012; Jakiela et al., 2013). In such systems, however, the cell suspension and reagents are handled by pumps and passed through the pumps directly, which not only introduces a significant risk of contamination by other microorganisms but also makes it difficult to clean the system. In addition, these systems don’t integrate their parts and didn’t demonstrate the ability to achieve sustained and reliable operation.

Here we report the development of an integrated platform referred to as a microbial microdroplet culture system (MMC), with which researchers can perform automated, high–throughput microbial cultivation and adaptive evolution in up to 200 replicate droplets of 2.00 μl volume. We integrated all the parts into a single chamber (Figure S1f), which resulted in a miniaturized, more reliable system. We also developed a new method for sample injection, which utilizes custom–designed reagent bottles that prevent contamination by other microorganisms and are much easier to be cleaned. In addition, we employed two Teflon tubes with good gas permeability, which enhances the gaseous exchange between droplets and external environment, to store and incubate droplets, allowing the microorganisms in the droplets to grow in a more preferred condition (such as higher concentration of dissolved oxygen) compared with cultivation on a chip. MMC allows automated, high–throughput cultivation of a variety of microorganisms with real–time monitoring of growth. With droplet manipulation on a chip, MMC also allows for continuous sub–cultivation, enabling the automated, high–throughput adaptive evolution of microorganisms to obtain strains that exhibit improved growth characteristics under certain stresses. In summary, MMC enables automated and sustained high–throughput cultivation and real–time monitoring of various microbial cells in droplets, and the system can also be used for adaptive evolution of microorganisms.

## MATERIALS AND METHODS

### 2.1. Materials

For our experiments, seven species of microbial cells were used: *E. coli* MG1655, *Lactobacillus plantarum* subsp. *plantarum* CICC 20418, *Corynebacterium glutamicum* ATCC 13032, *Saccharomyces cerevisiae* BY4741, *Pichia pastoris* Ppink– HC S–1, *Methylobacterium extroquens* AM1 (Liang et al., 2018), and a methanol– essential *E. coli* strain version 2.2 (MeSV2.2) (Meyer et al., 2018). Table S1 summarizes the culture medium used for each strain. Each strain was maintained in a strain–specific medium containing 20% glycerol (GENERAL–REAGENT, China) and frozen as pelleted cells at –80°C. Each strain was streaked on an agar plate prepared with the appropriate medium and incubated until single colonies appeared. Single colonies were picked and used to inoculate the appropriate liquid medium. Cells were then cultivated using the conditions shown in Table S2.

### 2.2. The working principles of MMC

MMC comprises four main parts: (1) pumps and valves, (2) reagent bottles, (3) a fluidic chip for droplet manipulation, and (4) a microbial cell cultivation tube (Figure 1a, refer to Figure S1 for photographs of the MMC components).

**Figure 1.**
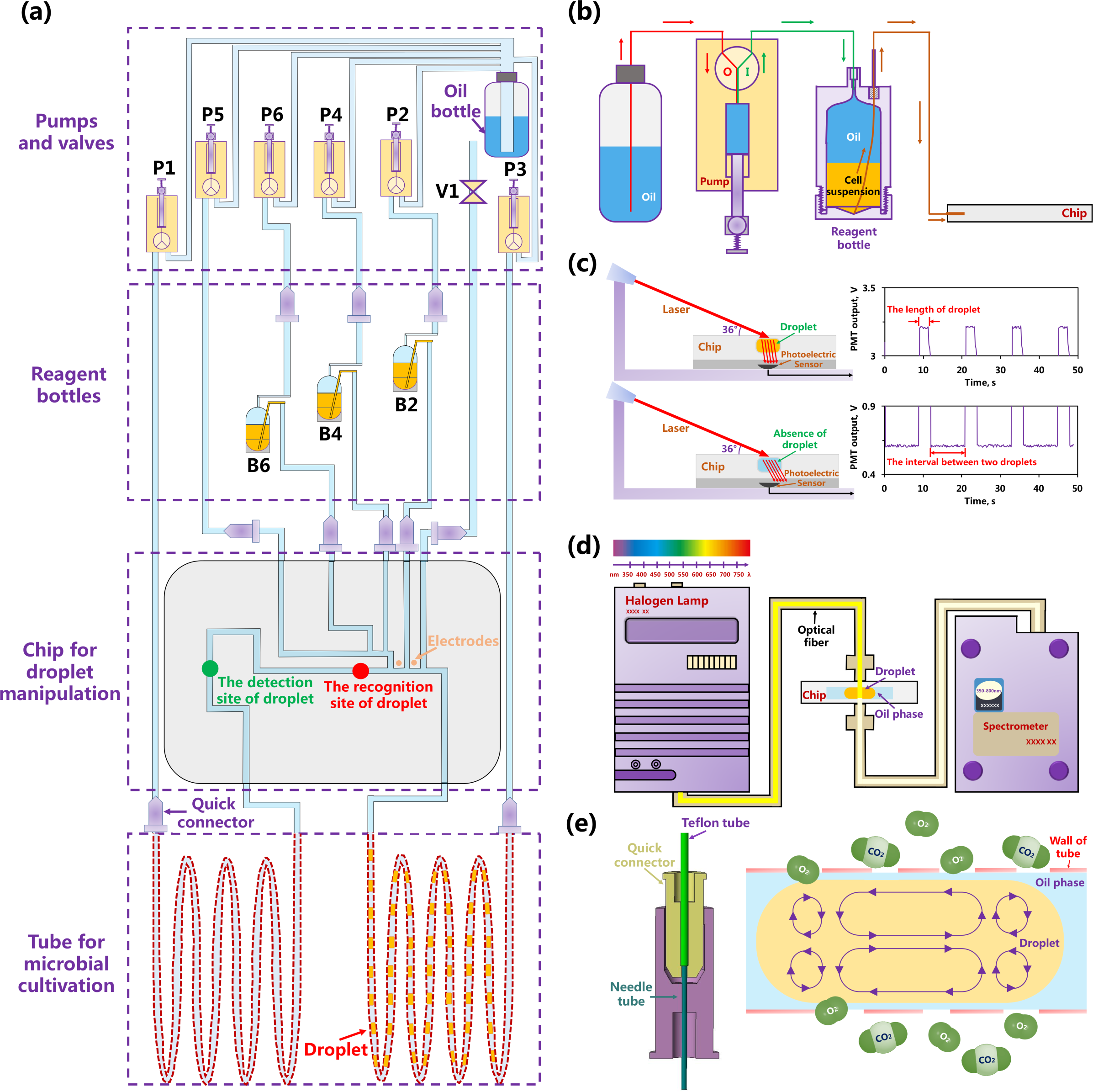
Schematic depiction of the MMC system. **(a)** The integrated framework for MMC. In the group of pumps and valves, there are six pumps (P1–6) and an electromagnetic valve (V1), which control the seven pipelines linked to the chip for droplet manipulation. In the group of reagent bottles, there are three special reagent bottles (B2, B4 and B6) containing the cell suspension (B2), culture medium (B4), and chemical factor (B6). The chip for droplet manipulation carries out the generation, cultivation, splitting, fusion, and sorting of droplets, which are identified and numbered at the recognition site of droplet (red dot). The optical density (OD) of each droplet is measured in real time at the detection site of droplet (green dot). The electrodes (orange dots) cause droplets to merge using an electrocoalescence mechanism. The tube for microbial cultivation is made of Teflon, which has good gas permeability. At certain pipeline joints, a quick connector is used for quick and easy connection of different pipelines. **(b)** Method for sample injection. In state O, the pump suctions oil from the oil bottle. In state I, the pump pushes the oil out, so the sample in the reagent bottle is pushed into the chip via the side tube. **(c)** Schematic of the recognition of droplets. The detection laser and the plane of the chip are at an angle of 36°. Owing to the difference in refractive index, when the droplet passes, the laser deviates more after passing through the droplet, then the photoelectric sensor below the chip receives more light and outputs a higher value of signal (>2 V). When the droplet is absent, the laser deviates less, and thus the photoelectric sensor receives less light and outputs a lower signal (<1 V). The length of the droplet and the interval between two adjacent droplets can be determined by the duration time of the signal. **(d)** Schematic of the monitoring of microorganisms in droplets. The halogen lamp, optical fiber, and spectrometer are arranged such that, when the droplet comes to the detection site, the OD (the wavelength range of detection is 350 nm ∼ 800 nm) inside the droplets is measured in real time. **(e)** Schematic of the structure of the quick connector and the gas exchange in droplets. Left: structure of a quick connector. Right: gas exchange in droplets. The tube for microbial cultivation (Teflon with good gas permeability) enables the exchange of gases inside and outside the droplets, thereby promoting the growth of microorganisms in the droplets.

#### 2.2.1. Pumps and Valves

MMC uses six pumps (Cavro Xcalibur Pump, TECAN, Switzerland) and an electromagnetic valve (Mrv–01–T02–K1.5–C–M01, Runze Fluid, China) controlled by a program created in–house using the programming application LabView. Each pump has three different connection states (Figure S2a) controlling sample injection and fluid flow in the system, and each pump regulates a pipeline. All pipelines are connected to an oil bottle at one end that supplies the carrier oil (mineral oil with 10 g/l Span 80, Sigma Aldrich, Germany) for the entire system. An electromagnetic valve controls the pipeline of waste discharge.

#### 2.2.2. Reagent bottles

Bespoke 15–ml bottles with a top tube and side tube were used as the reagent bottles (Figure S2b). These tubes are important for sample injection to prevent sample evaporation and contamination to achieve long–term automated continuous cultivation. Autoclaved (121°C, 15 min) reagent bottles are filled with 8–12 ml of oil via the side tube in a laminar flow cabinet. Then, 2–6 ml of sample is added via the same tube. Oil is then added to fill the remaining space in the bottle. The top tube is connected to a pump, and the side tube is connected to the chip. The sample, which occupies the lower part of the reagent bottle owing to its greater density, is pushed into the chip via the side tube when oil is pumped into the reagent bottle (Figure 1b).

#### 2.2.3. Droplet–manipulation fluidic chip

The droplet–manipulation fluidic chip, which is made of poly(methyl methacrylate), contains a channel that has an inner diameter of 1 mm. The chip has seven branches, each being connected to a pump, valve, reagent bottle, or microbial cell cultivation tube by an acrylonitrile butadiene styrene quick connector (Figure 1e).

The fluidic chip contains a recognition site of droplets, a detection site of droplets, and two electrodes for electrocoalescence. The recognition site of droplets consists of a laser (620 nm), photoelectric sensor (TSL12T, TAOS, USA), mechanical supports, and related circuits (Figure 1c). Each droplet passing through this site is numbered in sequence by the program for traceability. In addition, the distance between two adjacent droplets is adjusted and maintained by replenishing or draining the oil between them to prevent the fusion of droplets, ensuring the stability of the droplet sequence in the channel.

The detection site of droplets contains a halogen lamp (HL–2000, Choptics, China), optical fiber (Choptics), and a spectrometer (EQ–2000, Choptics) to monitor microbial growth in real time (Figure 1d). For our experiments, the system was set to measure the optical density at 600 nm (OD_600_) of each droplet over a 1–mm optical path when it passed through the detection site.

Finally, the pair of electrodes provide an alternating current electric field (1600V, 60 kHz) for electrocoalescence in which droplets are fused (Szymborski, et al., 2011).

#### 2.2.4. Microbial cell cultivation tube

Teflon microbial cell cultivation tubes (AF–2400, O.D. 1.67 mm, I.D. 1.07 mm, length 1.5 m) were used to store and incubate droplets. The high total surface area to volume ratio of the fluid increases the rate of mass transfer in the droplets, while the movement enhances convection, which in turn facilitates mixing in the droplets (Kaminski et al., 2016). In addition, gaseous exchange (Figure 1e) is enhanced by the gas permeability (Zhang et al., 2017a) of the Teflon AF–2400 tubes used for microbial cell cultivation and internal flow of droplets (Li et al., 2012).

### 2.3. On–chip droplets manipulation

Droplets can be manipulated in five different ways on the chip: (1) generation, (2) incubation, (3) splitting, (4) fusion, and (5) sorting (Figure 2).

**Figure 2.**
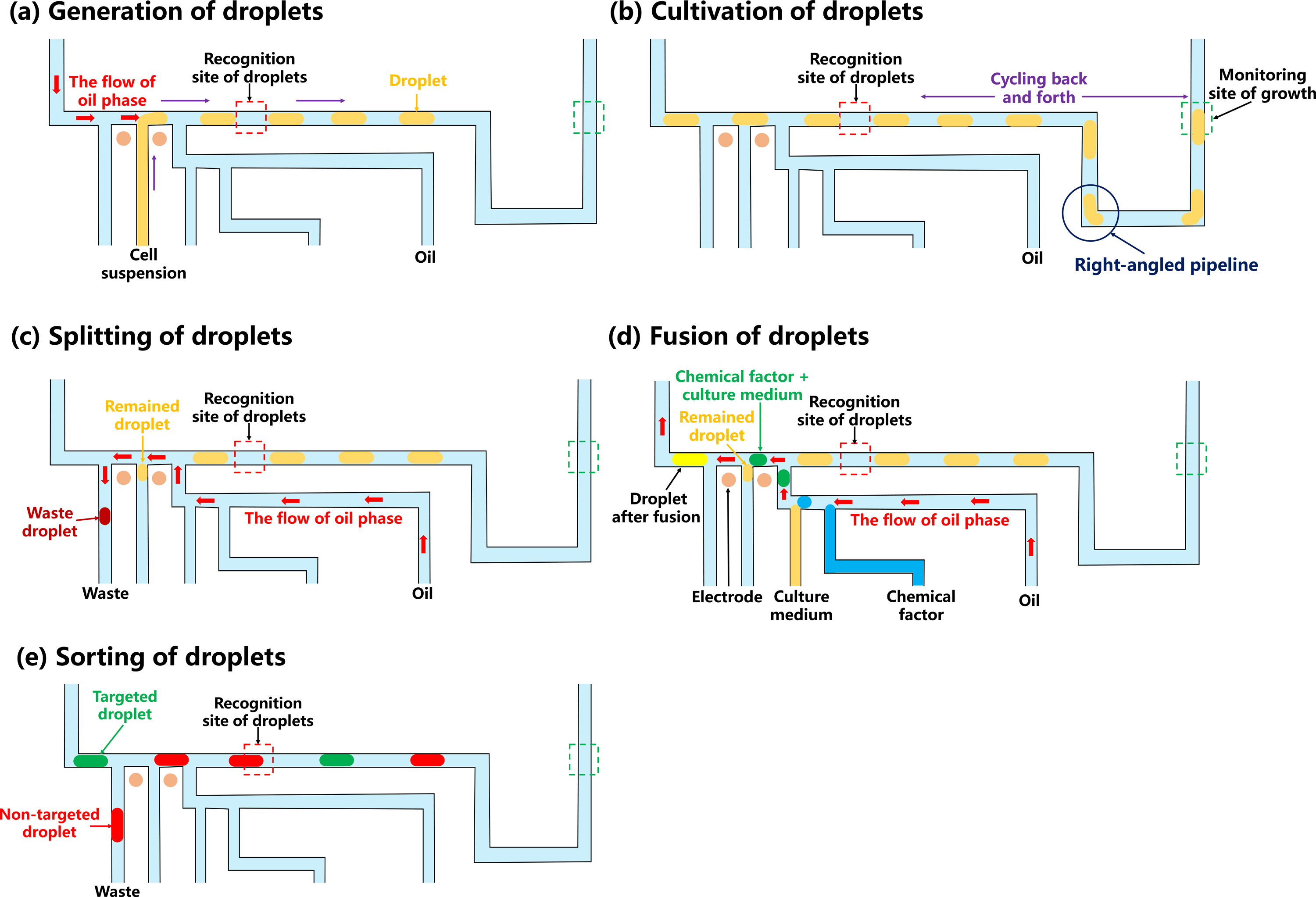
Schematic of the operations used to manipulate droplets on the chip. **(a)** Generation of droplets. Cell suspension from the reagent bottle is sheared at the T– junction by the oil phase coming from the vertical direction to form droplets. **(b)** Cultivation of droplets. After droplets are formed, they are cycled back and forth in the tube for microbial cultivation. Right–angled pipelines on the chip enhance the mixing of substances inside the droplets. At the monitoring site of growth, the OD is measured to monitor cell growth in real time. **(c)** Splitting of droplets. Droplets are sheared at the T–junction by the oil phase coming from the vertical direction to be split into the remaining droplet and a waste droplet. **(d)** Fusion of droplets. The remaining droplet is fused with a droplet containing fresh culture medium and chemical factor using an electrocoalescence mechanism and becomes a new droplet. In this system, the volume of the new droplet is the same as the droplet before splitting. **(e)** Sorting of droplets. The selected droplet proceeds to the tube for microbial cultivation, and the unselected droplet goes to the waste.

The MMC generates water–in–oil droplets, with their volume determined by the pumps and valves (Cedillo–Alcantar et al., 2019; Jakiela et al., 2013). For our experiments, the volume of each droplet generated was 2.00 μl, and the maximum number of droplets in each set of experiments was 200. The droplets are numbered using the program as they pass through the recognition site located after the droplet– generation site (Figure 2a, Video S1).

For incubation, the droplets are circulated in the droplet–manipulation fluidic chip and the microbial cell cultivation tube. The OD_600_ of droplets is measured as they pass through the detection site. The channel also contains some right–angled curves to enhance mixing by convection (Song et al., 2003; Song et al., 2006) (Figure 2b, Video S2).

Droplets splitting and fusion are two important aspects necessary to achieve continuous cultivation, which can conduct the operation of cell passage by splitting the droplets containing cells into two parts and then merging one of them with the droplets containing fresh medium (Figure 2c and 2d, Video S3). The fraction of splitting, *f*_*sp*_ is defined as

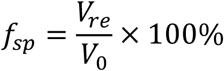

where *V*_*re*_ is the volume of the remaining droplet after being split and *V*_*0*_ is the initial volume of the droplet. By maintaining the volume of the droplet after fusion the same as the initial volume of droplet, the fraction of splitting, *f*_*sp*_, determines the concentration of the microbial cells present in the droplet after fusion, which can help us determine the concentration of inoculum when sub–cultivating cells. The fraction of splitting can be manually defined for the particular needs of each experiment. Droplets can be sorted and extracted using a droplet–numbering system in MMC (Figure 2e, Video S4).

### 2.4. Calibration of the measurement of OD_600_ in MMC and well plate

For our experiments, OD_600_ was used to determine the density of microbial cells in liquid culture medium. However, the Beer–Lambert law relating optical density to sample concentration (or cell density) is only applicable when the optical density of the sample solution falls within the linear range of the optical density vs. concentration calibration curve (Begot et al., 1996). Generally, before the OD_600_ of a microbial culture is measured, the culture is diluted to yield measurements that fall within the linear range of the spectrophotometer. However, manual dilution for spectrophotometer measurement is difficult in both the well plate and MMC formats. Therefore, it was necessary to plot calibration curves correlating OD_600_ values for each of these different formats to OD_600_ values obtained using a standard method. To generate these calibration curves, the OD_600_ values for various dilutions of each microbial species were first measured with a UV–Vis spectrophotometer (Ultrospec 3100 pro, GE, USA; linear range: 0.1–0.8). For high–density microbial cell suspensions, the culture was concentrated by centrifugation. To create high–density microbial cell suspensions, the cells were cultivated in a 100–ml shake flask using 20 ml of the appropriate liquid medium for a long enough time to reach the maximum cell density (for the time, refer to the growth curves of the shake flask in Figure 5). Then the cell suspension was transferred to a centrifuge tube and spun at 5000 rpm for 5 min at 4 °C. After centrifugation was completed, discarded a part of supernatant, and then resuspended the cells to obtain high–density microbial cell suspensions. Microbial cell suspensions of the same densities used for the UV–Vis measurements were then analyzed with a microplate reader (infinite M200 PRO, TECAN) and in the MMC system. Each measurement was performed in triplicate. The calibration curves for each strain were constructed by plotting the mean OD_600_ measured with the UV– Vis spectrophotometer against the mean OD_600_ measured with the microplate reader or in the MMC system (Figure 3 and S3).

**Figure 3.**
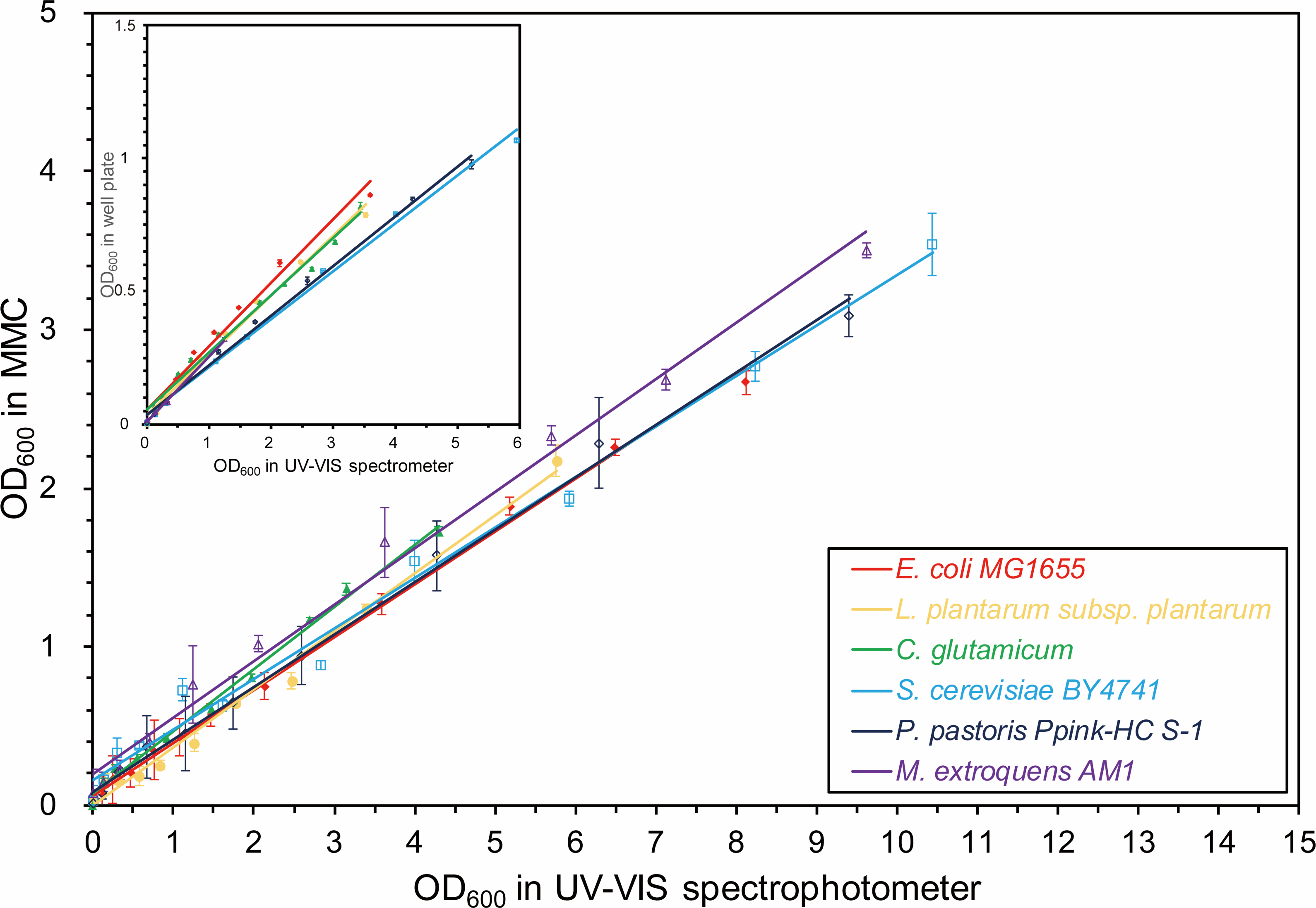
Calibration curves for OD_600_ measurements in well plates and MMC. In each situation, i.e., well plate (inset) or MMC, calibration curves for the various dilutions of six different trains are determined and plotted against measurements of the same dilutions made with a UV–Vis spectrophotometer. Each calibration curve is shown in Figure S3 in detail. The calibration curves of the six strains were quite similar for both the well plates and MMC. The linear range of the measurement of OD_600_ in MMC (0–11) is much greater than that in a well plate (0–6).

**Figure 4.**
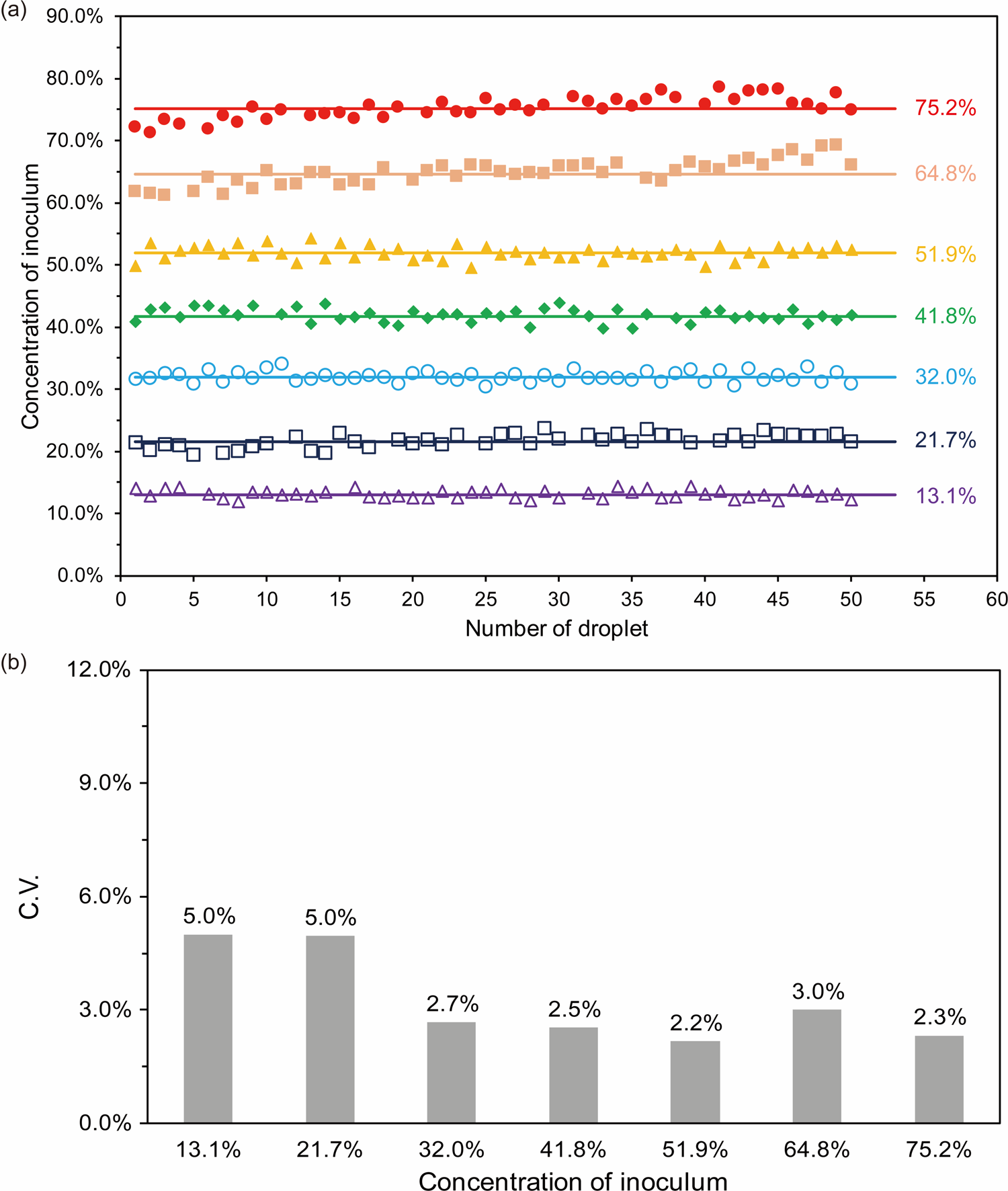
Characterization of the accuracy and reproducibility of inoculation in MMC. **(a)** Plot illustrating the measurement of the concentration of inoculum from seven experimental groups. The concentrations of inoculum in each experimental group were 13.1%, 21.7%, 32.0%, 41.8%, 51.9%, 64.8% and 75.2%. Fifty droplets were formed in each experimental group. **(b)** The coefficient of variation (C.V.) of 50 droplets in each experimental group in (a). The range of C.V. was 2.2% to 5.0%.

**Figure 5.**
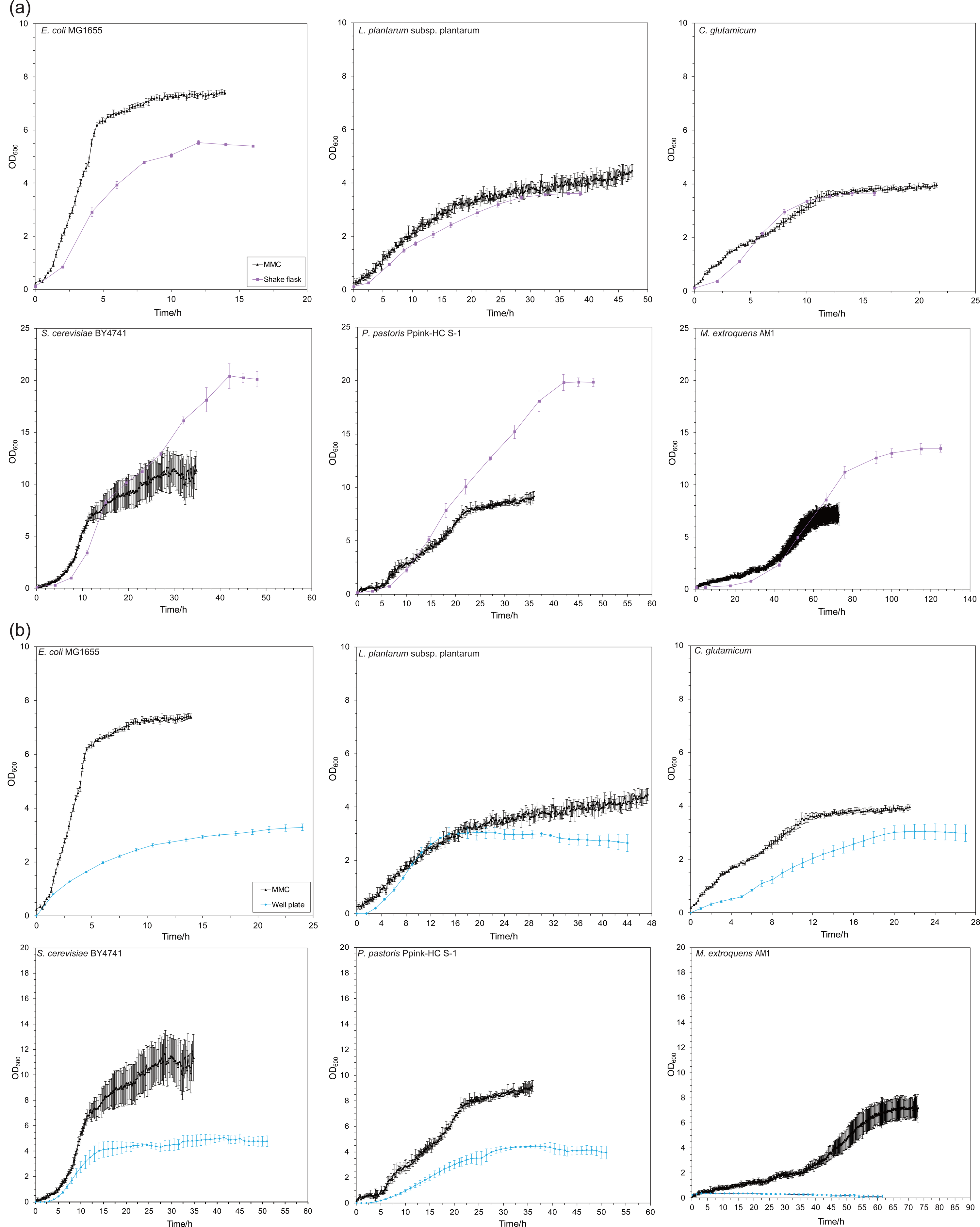
Growth curves for six strains cultivated in shake flasks, well plates, or MMC. For shake flasks and well plates, there were three parallel experimental groups. For MMC, 15 droplets were formed for every strain. All the growth curves are calibrated. **(a)** Comparison of MMC (black line) and shake flask (purple line). **(b)** Comparison of MMC (black line) and well plate (blue line).

### 2.5. Adaptive evolution of MeSV2.2 in MMC

MeSV2.2 was cultivated in a shake flask for 72 h using lysogeny broth culture medium. The culture medium was inoculated with MeSV2.2 suspension (2% of the total volume, Table S1) and cultivated in a shake flask for 5 h before seeding into MMC. Fifty droplets were generated, and the concentration of inoculum was set to 15% in each sub–cultivation. The duration of each sub–cultivation was 30 h. The culture medium was then replaced with fresh medium by droplet splitting and fusion.

## RESULTS

### 3.1. Calibration curve of OD_600_ values for MMC

We constructed OD_600_ calibration curves for six different microbial strains by plotting the OD_600_ values measured by UV–Vis spectrophotometry against OD_600_ values measured in both well–plate and MMC systems (Figure S3). Comparison among the calibration curves (Figure 3) revealed only a small effect of species difference on OD measurement in both well–plate and MMC formats, as the deviation among curves was minor. Notably, however, the shapes of all strains used for our experiment were rod–like or spherical, so this result might not be applicable to microbes of other shapes.

On the other hand, the linear range of the calibration curve varied among different instruments, a result of a difference in path length. The cuvette we used for UV–Vis spectrophotometry had a path length of 10 mm, and the linear OD_600_ range was 0.1– 0.8. For the well plate, the path length was 6 mm (the same as the height of the cell suspension in the well plate), and the linear OD_600_ range was 0–6. For MMC, the path length was 1 mm, and the linear OD_600_ range was 0–11. According to the Beer– Lambert Law, an increase in optical path length will increase the spectral noise of the system exponentially as a result of increased intensity of the strong absorption band (Inagaki et al., 2017). In turn, this increase in spectral noise increases the standard deviation of absorbance measurements, thereby narrowing the linear range of the absorbance curve.

### 3.2. Accuracy and reproducibility of inoculation

Automated, continuous cultivation in MMC is achieved by splitting and fusing droplets. Therefore, it is important to monitor the accuracy and reproducibility of the MMC inoculation process to gauge the impact of inoculation on parallelization. For ease of measurement, purple pigment and 10% sodium chloride solution were used to simulate microbial cells and culture medium. A series of solutions were prepared containing 10% sodium chloride and seven different concentrations of purple pigment (13.1%, 21.7%, 32.0%, 41.8%, 51.9%, 64.8% and 75.2%). For each sample, 50 droplets were generated. After the droplet–splitting process, the remaining droplets were inoculated into fresh 10% sodium chloride solution without purple pigment. The absorbance of the initial droplets and the fused droplets was measured at 600 nm, and the fraction of absorbance relative to that of the initial droplets was defined as the concentration of inoculum. The coefficient of variation of the concentration of inoculum relative to the targeted concentration varied between 2.2% and 5.0% for each sample (Figure 4), indicating that the inoculation process in MMC is reproducible. When the targeted concentration was 13.1% or 21.7%, the coefficient of variation was 5.00% and 4.96%, respectively, which were the highest among all samples (Figure 4b). This result is attributable to the higher precision requirement for the pump, both when segregating droplets into smaller fragments and droplet fusion. Given that the precision is consistent, the coefficient of variation will increase as the targeted concentration is decreased.

### 3.3. Characterization of the performance of microbial cultivation

To evaluate the performance of MMC with respect to microbial cultivation, six microbial–cell species were cultivated in shake flasks, well plates, and MMC, and the growth curves were plotted (Figure 5). For each type of microbial cell, a single colony growing on an agar plate was used to inoculate a 20–ml starter culture (see Table S1 for species–specific media and Table S2 for culture conditions) and cultivated in shake flasks. When the starter culture reached OD_600_ ≈ 0.1, aliquots (20 ml for a shake flask, 200 μl for a well and 2 μl for a droplet) were used to inoculate the various cultivation systems.

At early stages of cultivation, the growth rate of all microbial cells was higher in MMC compared with both shake flask and well–plate culture. With a larger total surface area to volume ratio, the droplet cultivation system had a higher mass–transfer rate, which enabled rapid gas exchange compared with other cultivation systems and thus provided more oxygen to support the growth of microbial cells while rapidly removing intracellular carbon dioxide. This property resulted in a higher initial growth rate when excess nutrients were present. At the middle and later stages of cultivation, when compared with shake–flask cultivation, *E. coli* MG1655 cultivated in the MMC grew much faster. In addition, *E. coli* MG1655 cultivated in the MMC had a higher–OD stationary phase. The cultivation performance of both *L. plantarum subsp. plantarum* (CICC 20418) and *C. glutamicum* (ATCC 13032) in the MMC was similar to that in the shake flask. However, the cultivation performance o *S. cerevisiae* BY4741, *P. pastoris* Ppink–HC S–1, and *M. extroquens* AM1 in MMC was inferior to that in the shake flask. For these three strains, the growth rate generally slowed when the OD_600_ reached ∼8, presumably because the droplet form inhibits cell survival at a high concentration of cells (Figure S4). In theory, when the cell concentration in the droplet is too high, the growth of these cells is inhibited because of several factors, such as space and nutrient limits, along with the accumulation of secondary metabolites in the droplets. The disadvantage in material transfer and the accumulation of secondary metabolites also explains the lower growth rate and final microbial cell density in well plates compared to MMC.

Furthermore, it is important to note that, although there was no growth of methanotrophic *M. extroquens* AM1 in the well–plate format, it exhibited a high initial growth rate in MMC. As a compartmentalized cultivation system, carrier oil in MMC insulates droplets from the environment, thus preventing the evaporation of any volatile compounds in the droplet (data not shown). During the cultivation of *M. extroquens* AM1, methanol was added to the culture medium to support growth. Methanol evaporated rapidly in the well–plate system, resulting in loss of the carbon source for *M. extroquens* AM1 and failure to grow. Although it is possible to add a layer of oil to the well–plate system to reduce evaporation, this approach renders further supplementation of methanol difficult and reduces oxygen diffusion into the culture medium and thus reducing the growth rate of *M. extroquens* AM1. These problems encountered in the well–plate system were not an issue with MMC.

### 3.4. Adaptive evolution of MeSV2.2 in MMC

Using microbial continuous sub–cultivation, we could conduct adaptive evolution to obtain microbial strains with desired properties. The engineering of synthetic methylotrophic microorganisms is of great significance for the conversion of one– carbon compounds in an industrial setting (Whitaker et al., 2015; Zhang et al., 2017b). Generally, engineering such synthetic microorganisms requires optimization in laboratory–based adaptive evolution experiments (Meyer et al., 2018). Here, we used MMC to perform adaptive evolution of methanotrophic MeSV2.2. As the highest density of MeSV2.2 we achieved was OD_600_ ≈ 0.3 after 40 h (Figure 6b and Figure S5), a mutant with higher growth rate and higher final cell density was desired. Considering the high evaporation rate of volatile compounds in well plates and the low throughput of shake–flask culture, MMC was clearly the superior option for adaptive evolution of MeSV2.2.

**Figure 6.**
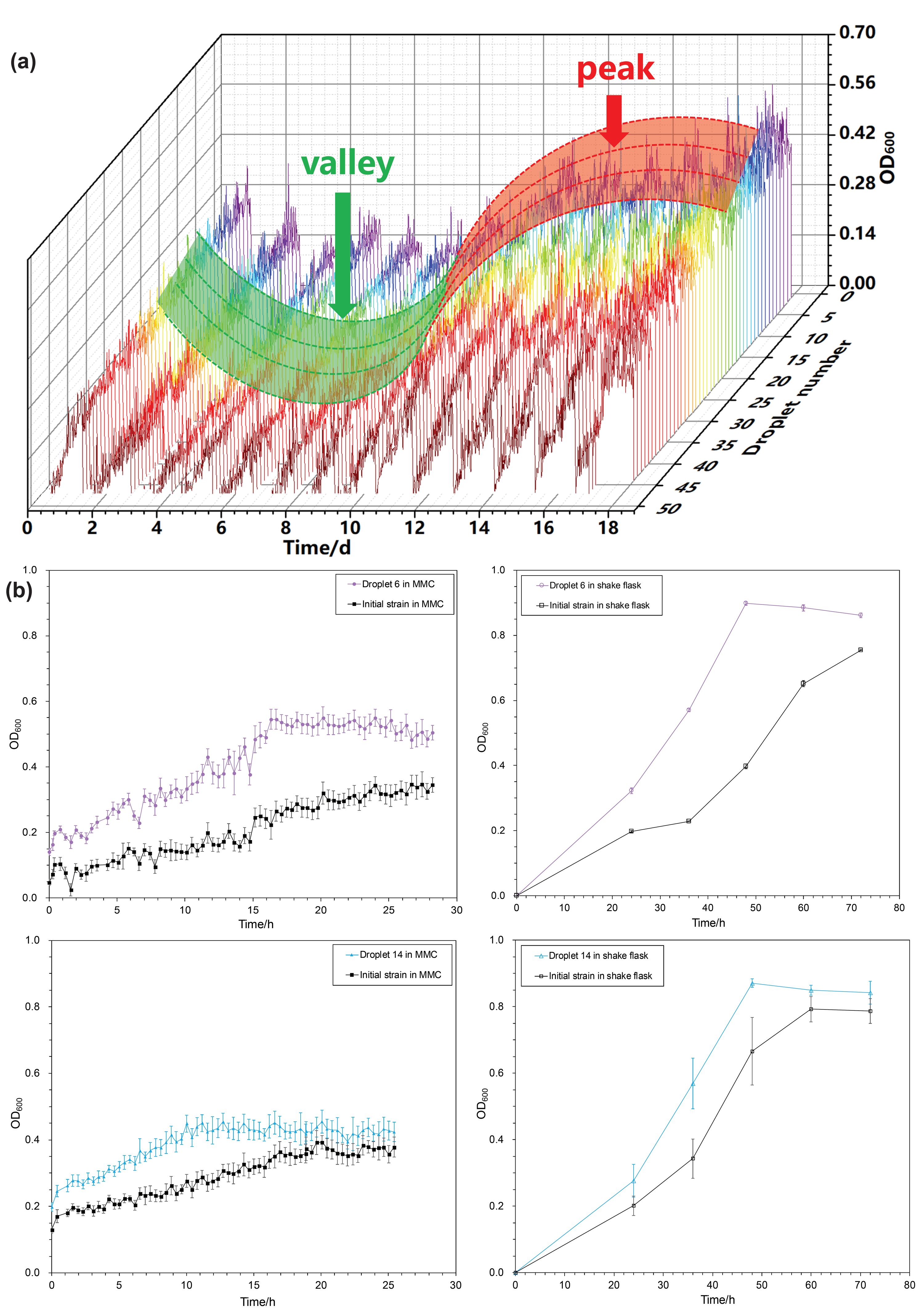
Results of the adaptive evolution of MeSV2.2 in MMC. **(a)** Growth curves for MeSV2.2 by continuous sub–cultivation. The highest points of the growth curves in each sub–cultivation first fell and then rose overall, resulting in the valley and peak in the figure. **(b)** Comparison of two selected strains and the initial strain. For the strains in droplets 6 and 14, whether they were cultivated in MMC or in shake flask, they all grew faster than the initial strain and had higher–OD stationary phases.

MeSV2.2 was cultivated in MMC for 18 days, and a growth curve for each droplet was plotted (Figure 6a). The peak and valley in each curve indicated the process of adaptive evolution of MeSV2.2. In MeSV2.2 culture medium, methanol was supplemented as a selection pressure for MeSV2.2. In the growth curves, inhibition of cell growth during the adaptation phase is indicated by the valley, whereas the higher growth rate of enriched cells adapted to methanol is indicated by the peak.

Eight droplets with high OD_600_ values after 18 days of cultivation were extracted for further investigation (droplets 2, 3, 6, 14, 20, 23, 28, 35). For the mutant strains in droplets 6 and 14, shake–flask cultivation yielded the highest growth rate (Figure S5). Furthermore, the growth rate of mutants in droplets 6 and 14 was compared with the parent strain in shake–flask and MMC cultivation (Figure 6b). The results revealed that the mutants in both droplets exhibited higher growth rates and had a higher cell density in the stationary phase when cultivated either in shake flasks or in MMC.

## DISCUSSION

We report the development of an automated, high–throughput microbial cultivation and adaptive evolution integrated platform called MMC that consists of four main parts, namely pumps and valves, reagent bottles, a droplet–manipulation fluidic chips, and microbial cell cultivation tubes, with novel designs (Figure 1), i.e., the sample– injection unit and quick connector. In similar work by Baraban et al. (2011) and Jakiela et al. (2013), the cell suspension and reagents flowed through the pumps directly and then were pushed into the chip. Because cleaning and sterilizing the pumps in this configuration is difficult, such a method not only renders the droplets highly susceptible to contamination by other microorganisms but also adds difficulty to system cleaning. In our novel method of sample injection, the cell suspension and reagents do not flow through the pumps, and the reagent bottles are easily cleaned and sterilized. The quick connectors connect modularized parts in the system, provide flexibility for parts replacement, and allow for straightforward system modification and customization.

The MMC has a large linear range of OD_600_ measurement (0 to ∼11), which eliminates the need for sample dilution prior to measurement via UV–Vis spectrophotometry. In addition, the small effect of different species on the OD_600_ calibration curves indicates the great reproducibility of the OD_600_ measurement system in MMC (Figures 3 and S3). These advantages enable continuous and in situ monitoring of microbial growth in MMC. Furthermore, it is possible to add fluorescence intensity–detection devices to MMC, enabling it to be used for analysis and sorting of microorganisms (Shen et al., 2019).

The concentration of inoculum is a crucial parameter in each sub–cultivation. In MMC, this parameter is determined by splitting and fusing droplets (Figure 2). When the concentration of inoculum was between 13.1% and 75.2%, the coefficient of variation (n = 50) was between 2.2% and 5.0% (Figure 4), indicating reproducible droplet manipulation in the MMC system. This reproducibly droplet manipulation system improves parallelization of each droplet, thereby maximizing the accuracy and validity of the results obtained.

The performance of microbial cultivation in MMC was investigated by growing six species of microbial strains and obtaining growth curves for each species in microliter–scale droplets (Figure 5). There are few reports on the growth of microorganisms in microscale droplets. Therefore, our results are important not only with respect to demonstrating the capability of long–term automatic cultivation in MMC but also for expanding the application of MMC, e.g., for laboratory–scale adaptive evolution. With the drawbacks of conventional cultivation techniques, i.e., a low–throughput and labor–intensive process, high reagent consumption of shake flasks, and poor mixing properties of well plates, MMC represents a great advantage in microbial cell cultivation. Use of a pump with greater power enabled manipulation of more droplets. The small volume of the reaction and the automated and reproducible operation of MMC greatly reduces reagent and labor costs and improves the parallelization of experiments. In a prior study, the maximum OD_600_ of the microbial cell cultures was less than 1 owing to gas–impermeability of the tube (Jakiela et al., 2013). The Teflon tube in MMC and the high mass transfer rate of the droplet system has improved microbial cell cultivation. In the present study, all six species of microbial strains were cultivated to high density (Figure 5). Although certain microbial strains, i.e., *S. cerevisiae* BY4741, *P. pastoris* Ppink–HC S–1 and *M. extroquens* AM1, had a lower growth rate in MMC compared to shake flask in the middle and late stages of cultivation, these parameters do not affect adaptive evolution of these strains in MMC as cultivation of microorganisms to a late stage is not required for adaptive evolution. Furthermore, the cultivation environment of MMC can be further improved by implementing a tube–in–tube reactor system (Zhang et al., 2017a). We can customize the gas atmosphere around the droplets, enabling microorganisms to grow under conditions of higher oxygen concentration or anaerobiosis according to experimental needs.

We also demonstrated adaptive evolution in MMC using the methanol–essential *E. coli* strain MeSV2.2 as a model. Generally, the equipment for maintaining continuous cultivation to sustain continuous evolution includes shake flasks, chemostat, an in– vial continuous cultivation system, a microfluidic–based continuous cultivation system, and a droplet–based continuous cultivation system (Tan et al., 2019b). MMC possesses many advantages when compared with such equipment, e.g., reduction in reagent consumption, high–throughput cultivation, and automated operation and high parallelization. In addition, modularization provides flexibility to the system. Owing to the great reliability of MMC, it is possible to conduct adaptive evolution experiments across several months.

Our results obtained with MMC suggests that this system can be adapted to many more applications in the future, such as the determination of the minimal inhibitory concentration of antibiotics (Andrews, 2001; Baraban et al., 2011; Kirchhoff et al., 2018), multi–factor optimization of microbial cultivation conditions (Ainala et al., 2016; Guo et al., 2009), and high–throughput cultivation and screening of single–cell droplets (Beneyton et al., 2017; Brouzes et al., 2009; Gruenberger et al., 2014). We expect that MMC will be used as a multi–functional integrated platform owing to its high throughput and automated operations.

## ACKNOWLEDGMENTS

This work was supported by the National Key Research and Development Program of China (2018YFA0901500), the National Key Scientific Instrument and Equipment Project of National Natural Science Foundation of China (21627812) and the Tsinghua University Initiative Scientific Research Program (20161080108).

## REFERENCES

Agresti, J. J., Antipov, E., Abate, A. R., Ahn, K., Rowat, A. C., Baret, J. C., Marquez, M., Klibanov, A. M., Griffiths, A. D., & Weitz, D. A. (2010). Ultrahigh–throughput screening in drop–based microfluidics for directed evolution. Proceedings of the National Academy of Sciences of the United States of America, 107(9), 4004–4009. doi: 10.1073/pnas.0910781107

Ainala, S. K., Seol, E., Kim, J. R., Ahn, K., & Park, S. (2016). Effect of culture medium on fermentative and CO–dependent H–2 production activity in Citrobacter amalonaticus Y19. International Journal of Hydrogen Energy, 41(16), 6734–6742. doi: 10.1016/j.ijhydene.2016.03.088

Akselband, Y., Cabral, C., Castor, T. P., Chikarmane, H. M., & McGrath, P. (2006). Enrichment of slow–growing marine microorganisms from mixed cultures using gel microdrop (GMD) growth assay and fluorescence–activated cell sorting. Journal of Experimental Marine Biology and Ecology, 329(2), 196–205. doi: 10.1016/j.jembe.2005.08.018

Andrews, J. M. (2001). Determination of minimum inhibitory concentrations. Journal of Antimicrobial Chemotherapy, 48, 5–16. doi: 10.1093/jac/48.suppl_1.5

Angell, J. B., Terry, S. C., & BarthA, P. W. (1983). Silicon Micromechanical Devices. Scientific American, 248(4), 44–55. doi: 10.1038/scientificamerican0483-44

Bachmann, H., Fischlechner, M., Rabbers, I., Barfa, N., dos Santos, F. B., Molenaar, D., & Teusink, B. (2013). Availability of public goods shapes the evolution of competing metabolic strategies. Proceedings of the National Academy of Sciences of the United States of America, 110(35), 14302–14307. doi: 10.1073/pnas.1308523110

Balagaddé, F. K., You, L. C., Hansen, C. L., Arnold, F. H., & Quake, S. R. (2005). Long–term monitoring of bacteria undergoing programmed population control in a microchemostat. Science, 309(5731), 137–140. doi: 10.1126/science.1109173

Baraban, L., Bertholle, F., Salverda, M. L. M., Bremond, N., Panizza, P., Baudry, J., de Visser, J. A. G. M., & Bibette, J. (2011). Millifluidic droplet analyser for microbiology. Lab on a Chip, 11(23), 4057–4062. doi: 10.1039/c1lc20545e

Begot, C., Desnier, I., Daudin, J. D., Labadie, J. C., & Lebert, A. (1996). Recommendations for calculating growth parameters by optical density measurements. Journal of Microbiological Methods, 25(3), 225–232. doi: 10.1016/0167-7012(95)00090-9

Beneyton, T., Thomas, S., Griffiths, A. D., Nicaud, J. M., Drevelle, A., & Rossignol, T. (2017). Droplet–based microfluidic high–throughput screening of heterologous enzymes secreted by the yeast Yarrowia lipolytica. Microbial Cell Factories, 16, 18. doi: 10.1186/s12934-017-0629-5

Brouzes, E., Medkova, M., Savenelli, N., Marran, D., Twardowski, M., Hutchison, J. B., Rothberg, J. M., Link, D. R., Perrimon, N., & Samuels, M. L. (2009). Droplet microfluidic technology for single–cell high–throughput screening. Proceedings of the National Academy of Sciences of the United States of America, 106(34), 14195–14200. doi: 10.1073/pnas.0903542106

Cedillo–Alcantar, D. F., Han, Y. D., Choi, J., Garcia–Cordero, J. L., & Revzin, A. (2019). Automated Droplet–Based Microfluidic Platform for Multiplexed Analysis of Biochemical Markers in Small Volumes. Analytical Chemistry, 91(8), 5133–5141. doi: 10.1021/acs.analchem.8b05689

Churski, K., Kaminski, T. S., Jakiela, S., Kamysz, W., Baranska–Rybak, W., Weibel, D. B., & Garstecki, P. (2012). Rapid screening of antibiotic toxicity in an automated microdroplet system. Lab on a Chip, 12(9), 1629–1637. doi: 10.1039/c2lc21284f

Cubas–Cano, E., Gonzalez–Fernandez, C., & Tomas–Pejo, E. (2019). Evolutionary engineering of Lactobacillus pentosus improves lactic acid productivity from xylose– rich media at low pH. Bioresource Technology, 288, 121540. doi: 10.1016/j.biortech.2019.121540

Dunham, M. J., Badrane, H., Ferea, T., Adams, J., Brown, P. O., Rosenzweig, F., & Botstein, D. (2002). Characteristic genome rearrangements in experimental evolution of Saccharomyces cerevisiae. Proceedings of the National Academy of Sciences of the United States of America, 99(25), 16144–16149. doi: 10.1073/pnas.242624799

Faassen, S. M., & Hitzmann, B. (2015). Fluorescence Spectroscopy and Chemometric Modeling for Bioprocess Monitoring. Sensors, 15(5), 10271–10291. doi: 10.3390/s150510271

Ferrer–Miralles, N., Domingo–Espin, J., Corchero, J. L., Vazquez, E., & Villaverde, A. (2009). Microbial factories for recombinant pharmaceuticals. Microbial Cell Factories, 8, 17. doi: 10.1186/1475-2859-8-17

Gao, X., Deng, L., Stack, G., Yu, H., Chen, X., Naito–Matsui, Y., Varki, A., & Galan, J. E. (2017). Evolution of host adaptation in the Salmonella typhoid toxin. Nature Microbiology, 2(12), 1592–1599. doi: 10.1038/s41564-017-0033-2

Grodrian, A., Metze, J., Henkel, T., Martin, K., Roth, M., & Kohler, J. M. (2004). Segmented flow generation by chip reactors for highly parallelized cell cultivation. Biosensors & Bioelectronics, 19(11), 1421–1428. doi: 10.1016/j.bios.2003.12.021

Gruenberger, A., Wiechert, W., & Kohlheyer, D. (2014). Single–cell microfluidics: opportunity for bioprocess development. Current Opinion in Biotechnology, 29, 15–23. doi: 10.1016/j.copbio.2014.02.008

Guo, W. Q., Ren, N. Q., Wang, X. J., Xiang, W. S., Ding, J., You, Y., & Liu, B. F. (2009). Optimization of culture conditions for hydrogen production by Ethanoligenens harbinense B49 using response surface methodology. Bioresource Technology, 100(3), 1192–1196. doi: 10.1016/j.biortech.2008.07.070

Hemmerich, J., Noack, S., Wiechert, W., & Oldiges, M. (2018). Microbioreactor Systems for Accelerated Bioprocess Development. Biotechnology Journal, 13(4), 1700141. doi: 10.1002/biot.201700141

Hong, K. K., & Nielsen, J. (2012). Metabolic engineering of Saccharomyces cerevisiae: a key cell factory platform for future biorefineries. Cellular and Molecular Life Sciences, 69(16), 2671–2690. doi: 10.1007/s00018-012-0945-1

Inagaki, T., Watanabe, T., & Tsuchikawa, S. (2017). The effect of path length, light intensity and co–added time on the detection limit associated with NIR spectroscopy of potassium hydrogen phthalate in aqueous solution. PLoS ONE, 12(5), e0176920. doi: 10.1371/journal.pone.0176920

Jakiela, S., Kaminski, T. S., Cybulski, O., Weibel, D. B., & Garstecki, P. (2013). Bacterial Growth and Adaptation in Microdroplet Chemostats. Angewandte Chemie International Edition, 52(34), 8908–8911. doi: 10.1002/biot.201700141

Kaminski, T. S., Scheler, O., & Garstecki, P. (2016). Droplet microfluidics for microbiology: techniques, applications and challenges. Lab on a Chip, 16(12), 2168–2187. doi: 10.1039/c6lc00367b

Kaushik, A. M., Hsieh, K. W., Chen, L. B., Shin, D. J., Liao, J. C., & Wang, T. H. (2017). Accelerating bacterial growth detection and antimicrobial susceptibility assessment in integrated picoliter droplet platform. Biosensors & Bioelectronics, 97, 260–266. doi: 10.1016/j.bios.2017.06.006

Kim, J., Shin, H., Park, H., Jung, H., Kim, J., Cho, S., Ryu, S., & Jeon, B. (2019). Microbiota Analysis for the Optimization of Campylobacter Isolation from Chicken Carcasses Using Selective Media. Frontiers in Microbiology, 10, 1381. doi: 10.3389/fmicb.2019.01381

Kirchhoff, J., Glaser, U., Bohnert, J. A., Pletz, M. W., Popp, J., & Neugebauer, U. (2018). Simple Ciprofloxacin Resistance Test and Determination of Minimal Inhibitory Concentration within 2 h Using Raman Spectroscopy. Analytical Chemistry, 90(3), 1811–1818. doi: 10.1021/acs.analchem.7b03800

Lee, K. S., Boccazzi, P., Sinskey, A. J., & Ram, R. J. (2011). Microfluidic chemostat and turbidostat with flow rate, oxygen, and temperature control for dynamic continuous culture. Lab on a Chip, 11(10), 1730–1739. doi: 10.1039/c1lc20019d

Li, P. F., Sun, H. B., Chen, Z., Li, Y., & Zhu, T. C. (2015). Construction of efficient xylose utilizing Pichia pastoris for industrial enzyme production. Microbial Cell Factories, 14, 22. doi: 10.1186/s12934-015-0206-8

Li, Z. H., Wu, J. K., Hu, G. Q., & Hu, G. H. (2012). Fluid flow in microfluidic chips (1st ed.). Beijing: Science Press.

Liang, W. F., Sun, M. Y., Cui, L. Y., Zhang, C., & Xing, X. H. (2018). Cre/loxP– Mediated Multicopy Integration of the Mevalonate Operon into the Genome of Methylobacterium extorquens AM1. Applied Biochemistry and Biotechnology, 185(3), 565–577. doi: 10.1007/s12010-017-2673-3

Lu, C., Lyumu, I., Xiaoling, D., Huan, D., & Zhangchao, D. (2018). Screening of copper–enriched microorganisms and optimization of culture conditions. New Biotechnology, 44, S82. doi: 10.1016/j.nbt.2018.05.915

Marose, S., Lindemann, C., Ulber, R., & Scheper, T. (1999). Optical sensor systems for bioprocess monitoring. Trends in Biotechnology, 17(1), 30–34. doi: 10.1016/S0167-7799(98)01247-5

MaymoGatell, X., Chien, Y. T., Gossett, J. M., & Zinder, S. H. (1997). Isolation of a bacterium that reductively dechlorinates tetrachloroethene to ethene. Science, 276(5318), 1568–1571. doi: 10.1126/science.276.5318.1568

Meyer, F., Keller, P., Hartl, J., Groninger, O. G., Kiefer, P., & Vorholt, J. A. (2018). Methanol–essential growth of Escherichia coli. Nature Communications, 9, 1508. doi: 10.1038/s41467-018-03937-y

Nautiyal, C. S. (1999). An efficient microbiological growth medium for screening phosphate solubilizing microorganisms. FEMS Microbiology Letters, 170, 265–270. doi: 10.1111/j.1574-6968.1999.tb13383.x

Nichols, D., Cahoon, N., Trakhtenberg, E. M., Pham, L., Mehta, A., Belanger, A., Kanigan, T., Lewis, K., & Epstein, S. S. (2010). Use of Ichip for High–Throughput In Situ Cultivation of “Uncultivable” Microbial Species. Applied and Environmental Microbiology, 76(8), 2445–2450. doi: 10.1128/AEM.01754-09

Nielsen, J., Larsson, C., van Maris, A., & Pronk, J. (1999). Metabolic engineering of yeast for production of fuels and chemicals. Current Opinion in Biotechnology, 24(3), 398–404. doi: 10.1016/j.copbio.2013.03.023

Ota, Y., Saito, K., Takagi, T., Matsukura, S., Morita, M., Tsuneda, S., & Noda, N. (2019). Fluorescent nucleic acid probe in droplets for bacterial sorting (FNAP–sort) as a high–throughput screening method for environmental bacteria with various growth rates. PLoS ONE, 14(4), e0214533. doi: 10.1371/journal.pone.0214533

Park, J., Kerner, A., Burns, M. A., & Lin, X. X. N. (2011). Microdroplet–Enabled Highly Parallel Co–Cultivation of Microbial Communities. PLoS ONE, 6(2), e17019. doi: 10.1371/journal.pone.0017019

Shen, Y. G., Yalikun, Y., & Tanaka, Y. (2019). Recent advances in microfluidic cell sorting systems. Sensors and Actuators B–chemical, 282, 268–281. doi: 10.1016/j.snb.2018.11.025

Sjostrom, S. L., Bai, Y. P., Huang, M. T., Liu, Z. H., Nielsen, J., Joensson, H. N., & Svahn, H. A. (2014). High–throughput screening for industrial enzyme production hosts by droplet microfluidics. Lab on a Chip, 14(4), 806–813. doi: 10.1039/c3lc51202a

Song, H., Bringer, M. R., Tice, J. D., Gerdts, C. J., & Ismagilov, R. F. (2003). Experimental test of scaling of mixing by chaotic advection in droplets moving through microfluidic channels. Applied Physics Letters, 83(22), 4664–4666. doi: 10.1063/1.1630378

Song, H., Chen, D. L., Tice, J. D., & Ismagilov, R. F. (2006). Reactions in droplets in microflulidic channels. Angewandte Chemie International Edition, 45(44), 7336–7356. doi: 10.1002/anie.200601554

Szymborski, T., Korczyk, P. M., Holyst, R., & Garstecki, P. (2011). Ionic polarization of liquid–liquid interfaces; dynamic control of the rate of electro–coalescence. Applied Physics Letters, 99(9), 094101. doi: 10.1063/1.3629783

Takahashi, C. N., Miller, A. W., Ekness, F., Dunham, M. J., & Klavins, E. (2015). A Low Cost, Customizable Turbidostat for Use in Synthetic Circuit Characterization. ACS Synthetic Biology, 4(1), 32–38. doi: 10.1021/sb500165g

Tan, Z. L., Ma, C. X., Xing, X. H., & Zhang, C. (2019a). Micro–cultivation system in microbiology: frontiers and prospects. Chinese Journal of Biotechnology, 35(7), 1151–1161. doi: 10.13345/j.cjb.190097

Tan, Z. L., Zheng, X., Wu, Y. N., Jian, X. J., Xing, X. H., & Zhang, C. (2019b). In vivo continuous evolution of metabolic pathways for chemical production. Microbial Cell Factories, 18, 82. doi: 10.1186/s12934-019-1132-y

Terry, S. C., Jerman, J. H., & Angell, J. B. (1979). Gas–chromatographic air analyzer fabricated on a silicon–wafer. IEEE Transactions on Electron Devices, 26(12), 1880–1886. doi: 10.1109/T-ED.1979.19791

Toprak, E., Veres, A., Michel, J. B., Chait, R., Hartl, D. L., & Kishony, R. (2012). Evolutionary paths to antibiotic resistance under dynamically sustained drug selection. Nature Genetics, 44(1), 101–105. doi: 10.1038/ng.1034

Toprak, E., Veres, A., Yildiz, S., Pedraza, J. M., Chait, R., Paulsson, J., & Kishony, R. (2013). Building a morbidostat: an automated continuous–culture device for studying bacterial drug resistance under dynamically sustained drug inhibition. Nature Protocols, 8(3), 555–567. doi: 10.1038/nprot.2013.021

Wadlin, J. K., Hanko, G., Stewart, R., Pape, J., & Nachamkin, I. (1999). Comparison of three commercial systems for identification of yeasts commonly isolated in the clinical microbiology laboratory. Journal of Clinical Microbiology, 37(6), 1967–1970. https://jcm.asm.org/content/37/6/1967.long

Whitaker, W. B., Sandoval, N. R., Bennett, R. K., Fast, A. G., & Papoutsakis, E. T. (2015). Synthetic methylotrophy: engineering the production of biofuels and chemicals based on the biology of aerobic methanol utilization. Current Opinion in Biotechnology, 33, 165–175. doi: 10.1016/j.copbio.2015.01.007

Wink, K., Mahler, L., Beulig, J. R., Piendl, S. K., Roth, M., & Belder, D. (2018). An integrated chip–mass spectrometry and epifluorescence approach for online monitoring of bioactive metabolites from incubated Actinobacteria in picoliter droplets. Analytical and Bioanalytical Chemistry, 410(29), 7679–7687. doi: 10.1007/s00216-018-1383-1

Wong, B. G., Mancuso, C. P., Kiriakov, S., Bashor, C. J., & Khalil, A. S. (2018). Precise, automated control of conditions for high–throughput growth of yeast and bacteria with eVOLVER. Nature Biotechnology, 36(7), 614–623. doi: 10.1038/nbt.4151

Zhang, C. S., & Xing, D. (2007). Miniaturized PCR chips for nucleic acid amplification and analysis: latest advances and future trends. Nucleic Acids Research, 35(13), 4223–4237. doi: 10.1093/nar/gkm389

Zhang, J. S., Teixeira, A. R., Zhang, H. M., & Jensen, K. F. (2017a). Automated in Situ Measurement of Gas Solubility in Liquids with a Simple Tube–in–Tube Reactor. Analytical Chemistry, 89(16), 8524–8530. doi: 10.1021/acs.analchem.7b02264

Zhang, W. M., Zhang, T., Wu, S. H., Wu, M. K., Xin, F. X., Dong, W. L., Ma, J. F., Zhang, M., & Jiang, M. (2017b). Guidance for engineering of synthetic methylotrophy based on methanol metabolism in methylotrophy. RSC Advances, 7(7), 4083–4091. doi: 10.1039/c6ra27038g

