## supplementary materials for "Microbial microdroplet culture system (MMC): an integrated platform for automated, high–throughput microbial cultivation and adaptive evolution"

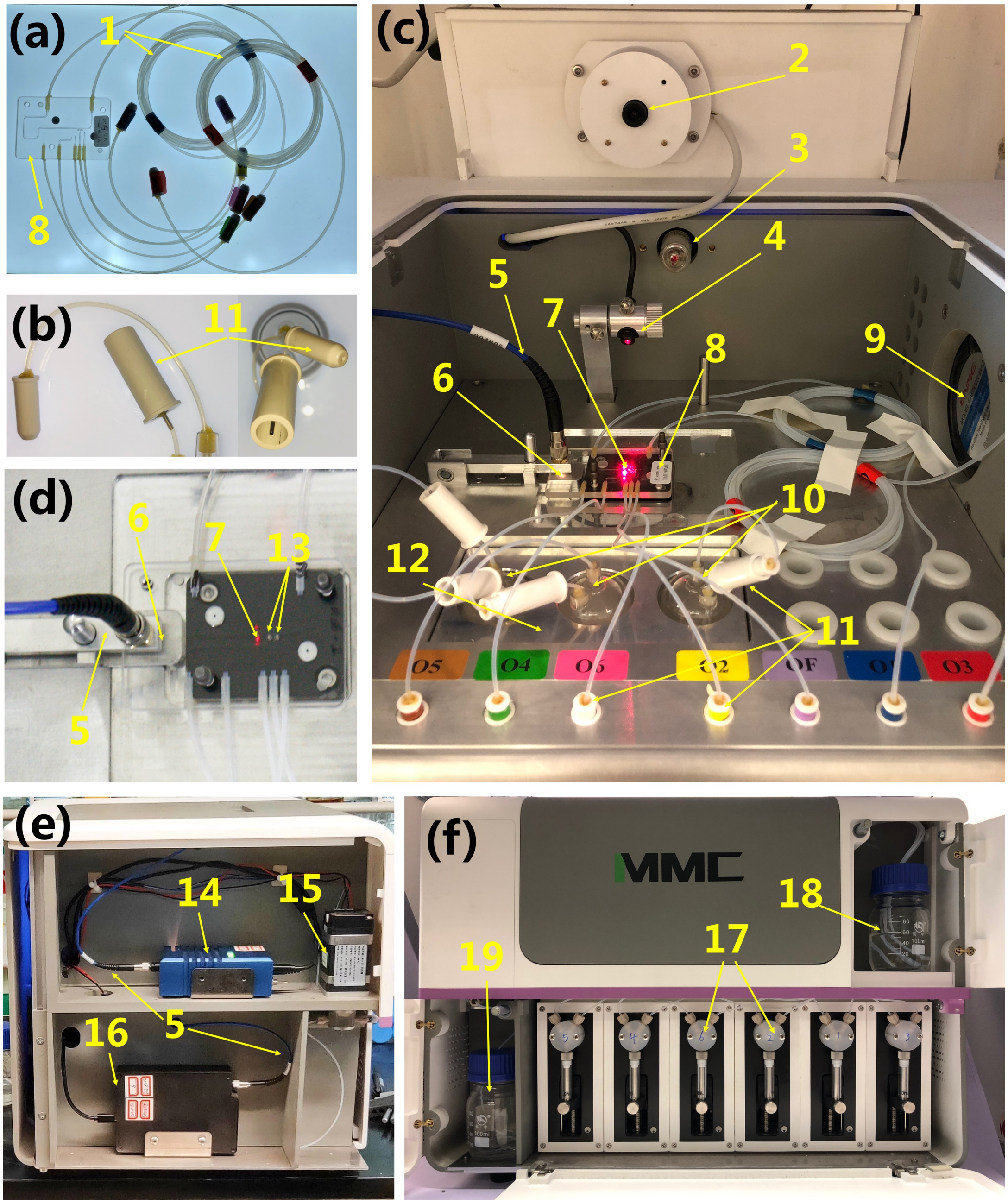


**Figure S1. Photographs of the microbial microdroplet culture system (MMC).** The system is 50 cm long, 40 cm wide, and 35 cm high. **(a)** Chip for droplet manipulation; **(b)** quick connector; **(c)** thermostatic chamber; **(d)** partial view of the chip in the thermostatic chamber; **(e)** inside of the left chamber of MMC; **(f)** front side of MMC.

1 — The tube for microbial cultivation.

2 — Camera for observing the formation and flow of droplets on the chip.

3 — UV lamp for sterilization. This lamp can be turned on in advance to sterilize the chip and tubes.

4 — Laser (620 nm) for droplet recognition.

5 — Optical fiber.

6 — Droplet-detection site.

7 — Droplet-recognition site (the photoelectric sensor is below).

8 — Chip for droplet manipulation.

9 — Heater for the thermostatic chamber. It can be used to maintain the temperature of microbial cultivation. The range of temperature that can be set is 3 ± 0.5°C to ~40 ± 0.5°C.

10 — Reagent bottles. These contain the cell suspension, culture medium, and chemical factors.

11 — Quick connector for rapid and easy connection of different pipelines.

12 — Metal bath. It can fix the reagent bottles and heat them to quickly raise the temperature of a reagent to the temperature of microbial cultivation.

13 — Electrodes, which provide an electric field for the electrocoalescence of droplets.

14 — Halogen lamp.

15 — Electromagnetic valve that controls the pipeline to the waste bottle.

16 — Spectrometer for the measurement of OD in droplets.

17 — Pumps that control sample injection and flow of fluids in the chip and tube.

18 — Oil bottle containing the oil used in the entire system.

19 — Waste bottle for collecting waste liquid.

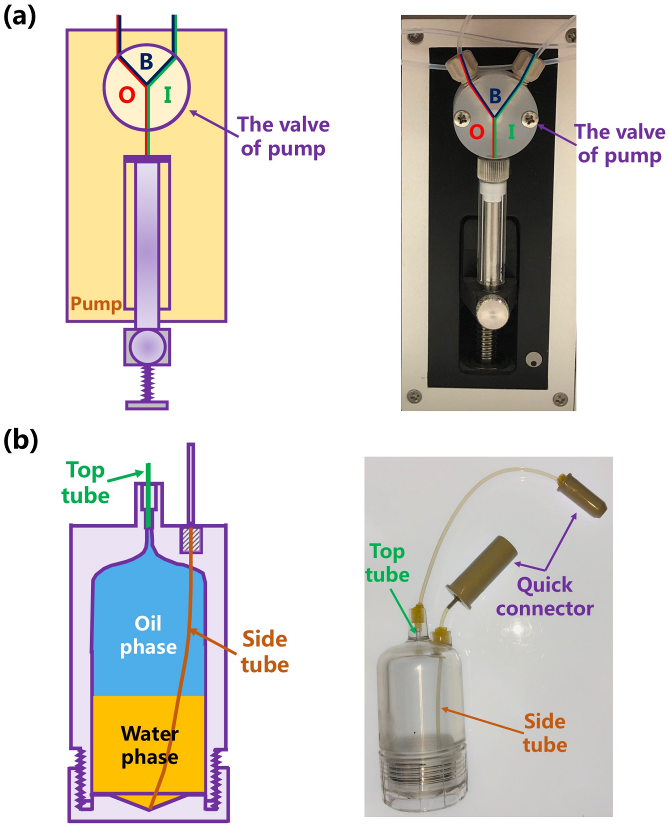


**Figure S2. Pump configurations and structure of the reagent bottles. (a)** Each pump has three different connection states. State O: the pump is connected to the oil bottle, and the pump can suction oil from the oil bottle in this state. State I: the pump is connected to the chip, and the pump can push the oil to the chip. State B: the chip is directly connected to the oil bottle, and the pipeline controlled by this pump is unimpeded. **(b)** The reagent bottle has a top tube and a side tube. The top tube only reaches the top of the bottle and is connected to the pump. The side tube reaches the bottom of the bottle and is connected to the chip.

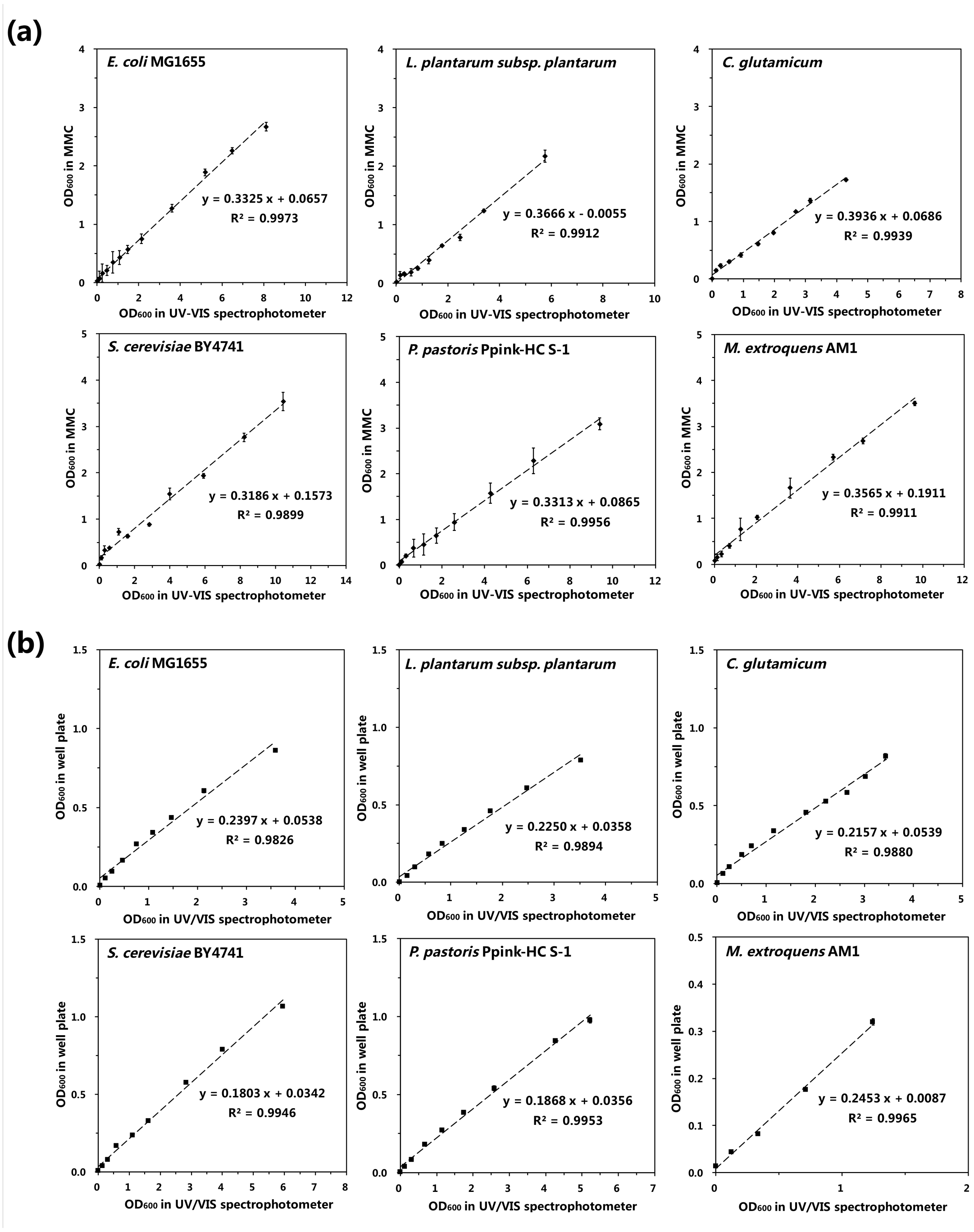


**Figure S3. Calibration curves of $OD_{600}$ measurements of the six different microbial strains in (a) MMC and (b) well plates.** For MMC, 10 droplets were formed for every point. For the well plate, three parallel wells were measured for every point.

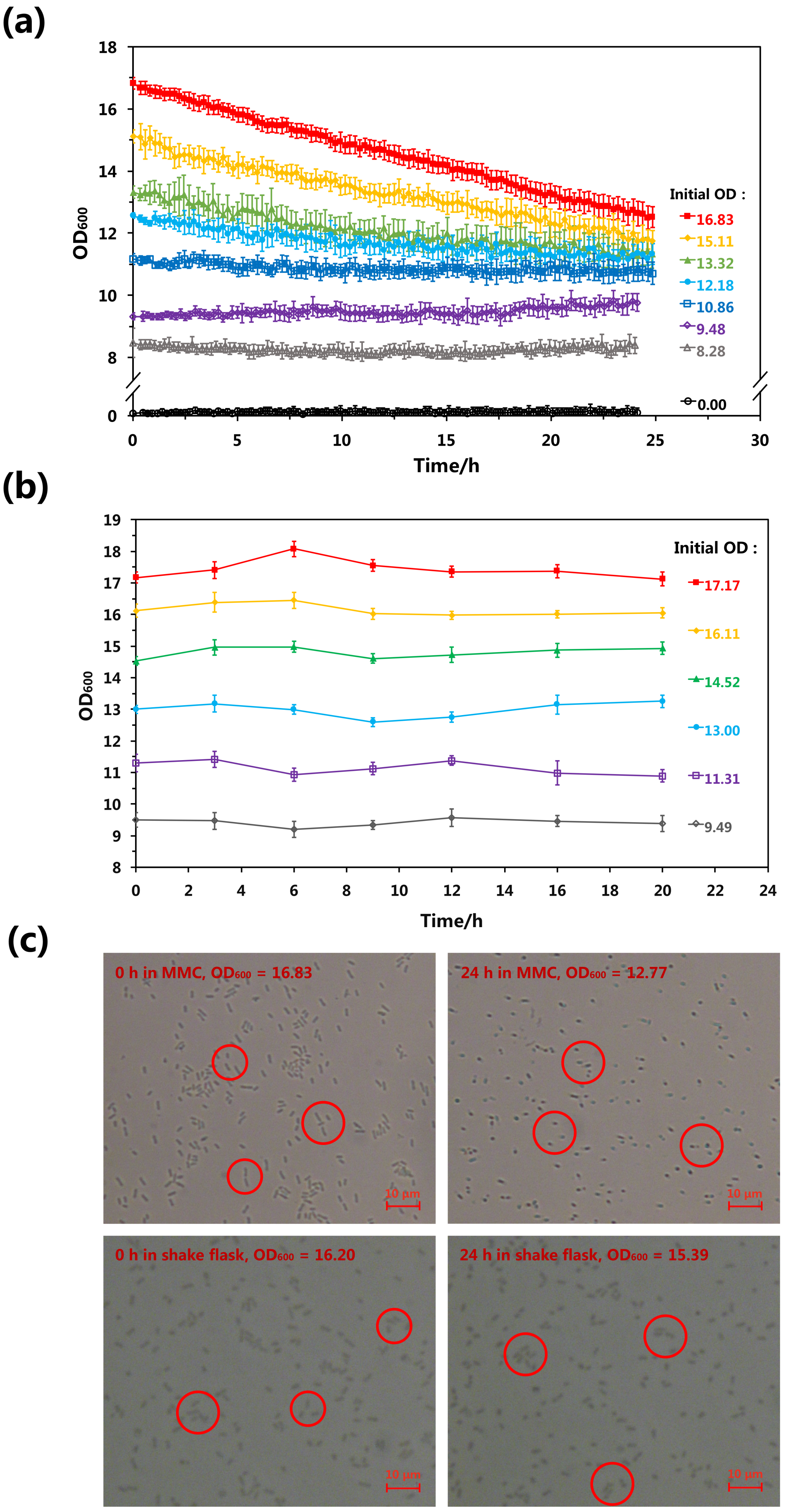
(a)
OD600
Time/h
Initial OD :
16.83
15.11
13.32
12.18
10.86
9.48
8.28
0.00
(b)
OD600
Time/h
Initial OD :
17.17
16.11
14.52
13.00
11.31
9.49
(c)
0 h in MMC, OD600 = 16.83
24 h in MMC, OD600 = 12.77
0 h in shake flask, OD600 = 16.20
24 h in shake flask, OD600 = 15.39
10 μm

**Figure S4. Change of *E. coli* MG1655 $OD_{600}$ over time at high concentration in shake flasks and MMC**. **(a)** $OD_{600}$ was measured over a period of 18 h in MMC (after calibration) and **(b)** shake flasks. *E. coli* MG1655 was cultivated in shake flasks for 18 h and then centrifuged at 5000 rpm for 5 min at 4 °C. To obtain high-concentration cell suspensions, the supernatant was removed and a small amount of phosphate-buffered saline (PBS) was added to resuspend the cells after centrifugation. In PBS, cells survive but cannot grow. The cells were then cultivated in a shake flask or MMC. **(c)** Photographs of *E. coli* MG1655 under microscope magnification. “0 h” and “24 h” refer to the initial state of cells in PBS and after 24 h of cultivation, respectively. In shake flasks, the high concentration of cells remained almost unchanged after 24 h. In MMC, however, the concentration of cells decreased over the same period. These decreases in concentration were most conspicuous in time courses beginning at higher initial OD values. The photographs indicate that, when cultivated in shake flasks, cell shape did not change substantially after 24 h of cultivation, i.e., cells remained completely rod-like. When cultivated in MMC, the shapes of cells became fragmentary after 24 h of cultivation. This indicates that the droplet structure may inhibit cell survival in a state of high concentration. This inhibition causes the cells to die and be degraded, resulting in a decrease in $OD_{600}$ and the fragmentary shapes of cells.

PBS: 8 g/l NaCl, 0.2 g/l KCl, 3.64 g/l $Na_2HPO_4 \cdot 12H_2O$ and 0.24 g/l $KH_2PO_4$, pH 7.4, stored at 4°C.

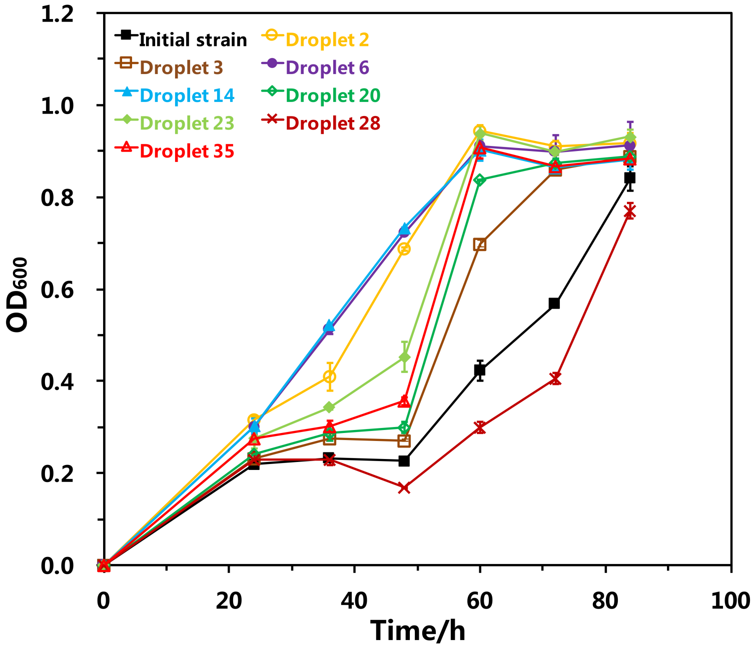


**Figure S5. Comparison of MeSV2.2 strains in eight selected droplets and the initial MeSV2.2 strain**. Growth curves were plotted for the MeSV2.2 strains in eight selected droplets as well as the initial MeSV2.2 strain in shake-flask culture. Three parallel groups were in the experiment. Strains in all droplets except droplet 28 grew faster than the initial strain. The strains in droplets 6 and 14 grew fastest.

**Table S1. Culture media used for seven microbial strains grown using MMC.** Unless specified, the solvent for all the solutions was deionized water. To prepare agar medium for plates, 15 g agarose was added per liter of medium. Media were autoclaved at 121°C for 15 min.

| Name of strain | Components of medium |
|---|---|
| *E. coli* MG1655 | Luria-Bertani culture medium:<br>10 g/l NaCl, 5 g/l yeast extract and 10 g/l tryptone |
| *Lactobacillus plantarum subsp. plantarum* CICC 20418 | 10 g/l glucose, 8 g/l tryptone, 2 g/l yeast extract, 17 g/l $CH_3COONa·3H_2O$, 0.2% (v/v) Solution A and 0.2% (v/v) Solution B, pH 6.8 ± 0.1.<br>Specifically,<br>Solution A: 50 g/l $K_2HPO_4$, 50 g/l $KH_2PO_4·3H_2O$, 0.2% (v/v) methylbenzene;<br>Solution B: 20 g/l $MgSO_4·7H_2O$, 1 g/l NaCl, 1 g/l $FeSO_4·7H_2O$, 1 g/l $MnSO_4·H_2O$. |
| *Corynebacterium glutamicum* ATCC 13032 | 10 g/l NaCl, 5 g/l yeast extract and 10 g/l tryptone |
| *Saccharomyces cerevisiae* BY4741 | 10 g/l yeast extract, 20 g/l peptone and 20 g/l Dextrose |
| *Pichia farinosa* Ppink-HC S-1 | 10 g/l yeast extract, 20 g/l peptone and 20 g/l Dextrose |
| *Methylobacterium extroquens* AM1 | 10% (v/v) Solution C, 0.055% (v/v) Solution D, 0.1% (v/v) Solution E, 1.5% (v/v) Solution F, 5% (v/v) methyl alcohol, 50 μg/ml kanamycin.<br>Specifically,<br>Solution C: 10 g/l $(NH_4)_2SO_4$, 4.5 g/l $MgSO_4·7H_2O$, 33 mg/l $CaCl_2·2H_2O$;<br>Solution D: 13.4 g/l $Na_3C_6H_3O_7·2H_2O$，345.04 mg/l $ZnSO_4·7H_2O$, 198 mg/l $MnCl_2·4H_2O$, 5 g/l $FeSO_4·7H_2O$, 2.46 g/l $(NH_4)_6Mo_7O_{24}·4H_2O$, 249.6 mg/l $CuSO_4·5H_2O$, 475.8 mg/l $CoCl_2·6H_2O$, 108.8 mg/l $Na_2WO_4·2H_2O$;<br>Solution E: 300 mg/l $H_3BO_3$;<br>Solution F: 402 g/l $Na_2HPO_4·12H_2O$, 130.5 g/l $KH_2PO_4$, pH = 6.8 ± 0.1. |
| MeSV2.2 | 6.78 g/l $Na_2HPO_4·12H_2O$, 3 g/l $KH_2PO_4$, 0.5 g/l NaCl, 1 g/l $NH_4Cl$, 0.34 g/l vitamin B1 (sterilized by filtration), 0.049 g/l $MgSO_4·7H_2O$, 1.5 mg/l $CaCl_2·2H_2O$.<br>Microelements: 0.5 mg/l $FeCl_3·6H_2O$, 0.09 mg/l $ZnSO_4·7H_2O$, 0.088 mg/l $CuSO_4·5H_2O$, 0.045 mg/l $MnCl_2$, 0.09 mg/l $CoCl_2·6H_2O$;<br>1.09 g/l gluconate, 500 mmol/l methanol, 0.1 mmol/l isopropyl-β-D-thiogalactopyranoside, 20 μg/ml streptomycin sulfate, 50 μg/ml kanamycin sulfate. |

**Table S2. Cultivation conditions for the six strains in shake flasks, well plates, and MMC.** For shake-flask cultures, strains were cultivated in a 100-ml shake flask using 20 ml of the corresponding liquid medium. For well-plate cultures, strains were cultivated in a 96-well plate (Costar, USA) using 200 μl of the corresponding liquid medium per well. To facilitate optical density measurements, each 96-well plate was placed in a microplate reader (infinite M200 PRO, TECAN, Switzerland) for cultivation.

| **Name of strain** | **Cultivation conditions** | | |
|---|---|---|---|
| | **Shake flask** | **Well plate (in microplate reader)** | **MMC** |
| *E. coli* MG1655 | 37°C, 200rpm | 37°C | 37°C |
| *Lactobacillus plantarum subsp. plantarum* CICC 20418 | 37°C, 200rpm | 37°C | 37°C |
| *Corynebacterium glutamicum* ATCC 13032 | 30°C, 220rpm | 30°C | 30°C |
| *Saccharomyces cerevisiae* BY4741 | 30°C, 220rpm | 30°C | 30°C |
| *Pichia farinosa* Ppink-HC S-1 | 30°C, 220rpm | 30°C | 30°C |
| *Methylobacterium extroquens* AM1 | 30°C, 170rpm | 30°C | 30°C |
